# Chemical representation standardization needed to generalize metabolic pathway involvement prediction across the Kyoto Encyclopedia of Genes and Genomes, Reactome, and MetaCyc knowledgebases

**DOI:** 10.1101/2025.04.02.646918

**Authors:** Erik D. Huckvale, Hunter N.B. Moseley

## Abstract

**Motivation:** Due to the utility of knowing the pathway involvement of compounds detected in biological experiments, knowledgebases such as the Kyoto Encyclopedia of Genes and Genomes (KEGG), Reactome, and MetaCyc have aggregated pathway annotations of compounds. However, these annotations are largely incomplete and are costly to obtain experimentally and curate from published scientific literature.

**Results:** We constructed a new dataset using compounds and their pathway annotations from KEGG, Reactome, and MetaCyc. Using this dataset, we trained and tested an extreme classification model that classifies 8,195 unique pathways based on compound chemical representations with a mean Matthews correlation coefficient (MCC) of 0.9036 ± 0.0033. During model evaluation, we discovered an inconsistency in chemical representations across knowledgebases, which was alleviated by standardizing the chemical representations using InChI (IUPAC International Chemical Identifier) canonicalization. Next, we compared the MCC between compounds and their cross-knowledgebase references. The non-standardized chemical representations had a huge 0.2687 drop in MCC while the standardized chemical representations only had a 0.0384 drop in MCC. Thus, standardizing chemical representation is an essential step when predicting on novel chemical representations.

**Availability and implementation:** All code and data for reproducing the results of this manuscript are available in the following figshare items:

Manuscript main results: https://doi.org/10.6084/m9.figshare.28701845

CV analysis of model and dataset of prior studies: https://doi.org/10.6084/m9.figshare.28701590

**Supplementary information:** ***<LINK TO SUPPLEMENTAL MATERIAL>***

## Introduction

Pathways are networks of interconnected chemical reactions within cells and organisms. If a chemical compound is involved as a product, reactant, or other small molecule participant of a chemical reaction, it is de facto associated with that reaction. And if a particular reaction takes place in a “pathway”, the compounds associated with that reaction are considered to be associated with that “pathway” (Voet, Voet and Pratt 2016; Berg *et al*. 2019; Nelson and Cox 2021). In this context, “pathway” can be a metabolic pathway, signaling pathway, biological process, disease process, or other biological concept with a graph-like representation of molecular interactions. When researchers detect various compounds within the biological samples of their experiments, it is highly useful to know which pathways the detected compounds are involved in, since that provides insight into the biological functions of the compound. This facilitates drug discovery, provides insight into causes and treatment of disease, and overall aids biological research. Because of this, the pathway associations of compounds are annotated in knowledgebases such as the Kyoto Encyclopedia of Genes and Genomes (KEGG) (Kanehisa and Goto 2000; Kanehisa 2019; Kanehisa *et al*. 2023), Reactome (Milacic *et al*. 2024), and MetaCyc (Caspi *et al*. 2020). However, these knowledgebases are incomplete, as there are many compounds without any pathway annotations, and determining the pathway involvement experimentally is time consuming and costly.

For this reason, several prior studies have prototyped machine learning models that predict the pathway involvement based on a compound’s chemical representation with varying levels of performance. Attempts to predict the pathway involvement of compounds most notably began with the work of Hu et al where the information of chemical interactions was used to predict 11 level 2 metabolic pathway categories found in KEGG (Hu *et al*. 2011). Building off of the work of Hu et al, Baranwal et al created a dataset representing compounds in the SMILES format (Weininger 1988) along with their mapping to one or more of the 11 KEGG level 2 metabolic pathway categories (Baranwal *et al*. 2020). Baranwal et al trained a multi-output graph neural network (Asif *et al*. 2021) with 11 outputs, one for each pathway category, where compounds were represented as a graph and information about their molecular structure was used to predict their pathway involvement. Yang et al (Yang *et al*. 2020) and Du et al (Du *et al*. 2022) later proposed different variants of graph neural networks to predict these same pathway categories using the same dataset. Huckvale and Moseley discovered that the results of the models trained on this initial dataset were invalid (Huckvale and Moseley 2024a) due to exact duplicate samples within the dataset, leading to data leakage and an overoptimistic measure of model performance (Yang *et al*. 2022). As a result, Baranwal et al published a corrected paper with the duplicate samples removed from the dataset (Baranwal *et al*. 2024). All studies published before September 21, 2024 used either a multiclassifier or a set of binary classifiers implementing a one-vs-rest classification approach and only predicted 11 or 12 level 2 metabolic pathways defined in KEGG. Since Baranwal et al met proper standards of scientific computational reproducibility by providing their code and data, we were able to train their model over 50 CV iterations and calculate MCC, resulting in a mean MCC of 0.7642 and standard deviation of 0.0137 (Table S1), providing representative performance of models generated prior to September 21, 2024. Moreover, it is important that model performance be reported in MCC due to the high imbalance in the training and testing datasets. With high imbalance, MCC has an advantage over the F1-score that ignores true negatives and major advantage over accuracy that ignores false positives and false negatives within the numerator (Cao, Chicco and Hoffman 2020; Chicco and Jurman 2020).

KEGG pathways are organized in a hierarchical fashion where there are seven top level (level 1) pathway categories within which there are second level pathway categories and then on the third level, we see individual pathways (KEGG Pathway Browser). The 11 outputs of these past models specifically were predicting the second level pathway categories under the ‘Metabolism’ top level category. While these initial models were instrumental in demonstrating the capability of predicting pathway involvement based on information of a compound’s molecular structure, the reality is that there are far more pathways that are of biological interest. KEGG alone has over 500 pathways defined (KEGG Pathway Browser). Meanwhile, Reactome and MetaCyc both have thousands of pathways defined (Reactome Pathway Browser; MetaCyc Pathway Browser). Therefore, this is not a simple multi-output problem but rather an extreme classification problem (Bengio et al. 2019)(Varma 2019) with thousands of different classes. One could train a multi-output model with thousands of outputs, but it is well known that as the number of classes increases while the dataset size remains the same, the more challenging it is to accurately predict the increasing number of classes (Moral, Nowaczyk and Pashami 2022). Alternatively, a separate binary classifier could be trained for each class, but in the case of pathway prediction, there are several small pathways with very few associated compounds. This results in many more negative entries than positive entries and the very high class imbalance greatly reduces model performance (Guo *et al*. 2008).

Huckvale and Moseley resolved the extreme classification problem with metabolic pathway prediction by developing a machine learning pipeline that cross joins compound features with features representing a pathway, training just a single binary classifier that predicts whether the given compound is associated with the given pathway (Huckvale and Moseley 2024b). With this technique, rather than the limited data set size (number of compounds only) needing to be shared amongst thousands of classes in a multi-output model, the dataset size increases, being multiplied by the number of classes (pathways), and only a single output is necessary. This is because rather than a dataset entry being defined as a compound, it is defined as a compound-pathway pair. This reformulation of the metabolic pathway prediction problem demonstrated that a model can be trained in a computationally practical manner while predicting an indefinite number of pathways with sufficient performance. Firstly, Huckvale and Moseley demonstrated that not only can 12 level 2 metabolic pathways be effectively predicted (we included the poorly performing pathway that everyone else left out) (Huckvale and Moseley 2024b), but also on 172 level 3 pathways (Huckvale and Moseley 2024c). This followed with predicting all 502 pathways defined in KEGG using a dataset with 6485 compounds. Going beyond KEGG, Huckvale and Moseley later demonstrated that models can effectively be trained to predict all 3985 Reactome pathways (Huckvale and Moseley 2025) and all 4055 MetaCyc pathways (Huckvale and Moseley 2024d). In addition, these studies demonstrated that training on all the pathways together with a single extreme classification model resulted in significant transfer learning between pathway classes that greatly improves the prediction of pathways compared to training a separate model per pathway in classic one-vs-rest approaches.

To handle this high number of pathway classes, a graph neural network, such as those that were trained in prior studies, was not immediately appropriate since Huckvale and Moseley entirely reformulated the problem to be capable of extreme classification. This includes the introduction of pathway features, which cannot feasibly be represented as graphs. A multi-layer perceptron (Bisong 2019), as compared to the graph neural networks used previously, is more practical for this extreme classification problem given implementation complication and hardware constraints. To predict the pathway involvement using a multi-layer perceptron (MLP), a vector representation of both compounds and pathways was needed. The compound vector representation was made possible by the work of Jin et al (Jin, Mitchell and Moseley 2020; Jin and Moseley 2021, 2023) in developing a graph-based atom coloring technique where the atoms of the compound are “colored” by the chemical substructure surrounding each atom. The atom coloring features for a compound are the counts of the atom colors present in the compound. The pathway features are likewise constructed by aggregating the compound features of the compounds associated with the pathway (Huckvale and Moseley 2024b). We continue to demonstrate in this work that a MLP is more than sufficient for this extreme classification task.

With models being able to effectively predict the pathways annotated in these three major knowledgebases, an intuitive hypothesis is that the mean model performance and model robustness can be further improved by training a model on a dataset constructed from compounds and pathways in KEGG, Reactome, and MetaCyc combined. We will refer to this as the KEGG+Reactome+MetaCyc dataset. However, the challenge with combining knowledgebases is their molfiles (Dalby *et al*. 1992) have inconsistent chemical representations. This impacts both the way that the compounds are represented in the compound features as well as the pathway features which are derived from the compound features. We demonstrate in this work that these chemical representation inconsistencies confuse the model. By standardizing with InChI canonicalization (Heller *et al*. 2013, 2015; Goodman *et al*. 2021), we make the chemical representations and therefore the input features consistent, further improving the predictive performance of all pathways across all three knowledgebases. This is similar to standardization methods used by PubChem; however, PubChem has different tautomeric preferences than InChI canonicalization (Hähnke, Kim and Bolton 2018). We also demonstrate that the InChI-based standardization greatly improves the generalizability of the model, enabling the effective prediction of pathway involvement of novel chemical representations.

## Materials and methods

### Creating the initial dataset

To create the KEGG+Reactome+MetaCyc dataset, we downloaded the compound molfiles from the KEGG, Reactome, and MetaCyc knowledgebases along with their pathway annotations. We used the kegg-pull Python package (Huckvale and Moseley 2023) to download the KEGG data (Huckvale and Moseley 2024e) and we downloaded the Reactome and MetaCyc data using their own web APIs directly (Huckvale and Moseley 2024d, 2025). Using the md-harmonize Python package (Jin and Moseley 2023), we generated the atom colors for each compound from its molfile and generated the corresponding atom color feature vector for each compound from the counts of of the atom colors present in the compound. The pathway feature vectors were sums of the feature vectors of the compounds associated with each pathway. So while the compound features are the counts of the atom colors present in each compound, the pathway features are the counts of the atom colors across all compounds within each pathway. Both the compound and pathway features were de-duplicated feature-wise and entry-wise, meaning any duplicate features were removed and any compound or pathway feature vectors with identical atom color counts were removed. In the case of duplicate pathway or compound feature vectors, the remaining unique feature vectors represented one or more compounds or pathways rather than strictly one. Since compounds feature vectors potentially represent more than one compound, the pathway annotations of all compounds that it represents are combined. Both the compound and pathway features are additionally normalized using softmax entry-wise and min-max scaling feature-wise (Huckvale and Moseley 2024b).

The pathway annotations served as the labels for the machine learning models such that we cross-joined the compound feature vectors to pathway feature vectors such that each entry in the dataset was a compound-pathway pair and the label was a binary value indicating whether the given compound is associated with the given pathway (Huckvale and Moseley 2024b). However, since the compounds and pathways came from different knowledgebases, only the compounds and pathways from the same knowledgebase are cross-joined, creating a block diagonalization by knowledgebase in the final matrix of entries. This reduces the number of negative labels, since the compounds from one knowledgebase are necessarily not associated with the pathways of a different knowledgebase.

Table 1 shows descriptive statistics of the dataset constructed from the combined data of KEGG, Reactome, and MetaCyc. There are more compound features and more pathway features, since both were generated from a larger statistical sample of molfiles. While one may expect that the number of compounds and the number of pathways in the KEGG+Reactome+MetaCyc dataset would be the sum of that of the three knowledgebases, this is not the case due to the de-duplication of the feature vectors, indicating that there were duplicate entries across the knowledgebases. The 16,660 unique compound feature vectors represent 18,554 compound entries across the knowledgebases. Likewise, 8,195 unique pathway feature vectors represent 22,504 pathway entries across the knowledgebases. However, a large number of completely duplicate pathways are present in Reactome, which is why the ratio of pathway entries to unique pathway feature vectors is above 2.5.

**Table 1.** Description of the combined KEGG+Reactome+MetaCyc dataset compared to that of prior studies.

| **Dataset** | **#Compound features** | **#Pathway features** | **#Unique Compounds** | **#Unique Pathways** | **#Entries** | **Reference** |
| --- | --- | --- | --- | --- | --- | --- |
| KEGG+Reactome+MetaCyc | 34,474 | 27,208 | 16,640 | 8,195 | 50,127,958 | Current study |
| KEGG | 16,509 | 11,321 | 6,485 | 502 | 3,255,470 | (Huckvale and Moseley 2024e) |
| Reactome | 6,187 | 5,386 | 1,976 | 3,985 | 7,874,360 | (Huckvale and Moseley 2025) |
| MetaCyc | 19,081 | 15,349 | 9,847 | 4,055 | 39,929,585 | (Huckvale and Moseley 2024d) |

To evaluate model performance, we performed 100 cross validation (CV) iterations where we created the train/test splits in a stratified manner (Verstraeten and Van den Poel 2006). Each CV iteration is a separate 10-fold cross-validation, but with only one CV fold tested. On each CV iteration, the train set had its positive entries duplicated until the number of positive samples roughly equaled the number of negative entries. While this is not valid to do in the test set, it greatly helped with the class imbalance during training. The resulting train set was then used to train a multi-layer perceptron model, with hyperparameters tuned using the Optuna Python package (Akiba *et al*. 2019). Once the model was trained, it predicted on the test set, collecting the number of true positives (TP), true negatives (TN), false positives (FP), and false negatives (FN). This enabled calculating metrics on each CV iteration including the Matthew’s correlation coefficient (MCC) (Cao, Chicco and Hoffman 2020; Chicco and Jurman 2020), F1-score, precision, recall, and accuracy. We counted the TP, TN, FP, and FN for all entries in the test set in order to calculate the overall mean, median, and standard deviation MCC across the CV iterations. And we counted the same per entry such that we could determine the per-knowledgebase MCC by counting the TP, TN, FP, FN of entries belonging to each of the three knowledgebases.

To evaluate the performance when the model is trained on the entries of one knowledgebase and pathways predicted on another knowledgebase, we split the KEGG+Reactome+MetaCyc dataset by knowledgebase, having a separate KEGG, Reactome, and MetaCyc dataset with the same input features. For each knowledgebase, we trained a model, using the same hyperparameters as the CV analysis, on the compounds and pathways of that knowledgebase and used the resulting model to predict the pathways of the compounds in the other two knowledgebases. We calculated the MCC of the predictions for each combination of training knowledgebase and prediction knowledgebase.

### Cross-reference analysis

To determine how well the model generalizes on different chemical representations from different knowledgebases, we determined how consistently the model predicts between pairs of compounds of the same identity cross-referenced between knownledgebases. We did this by taking molfiles of compounds along with their pathway annotations in one knowledgebase and obtaining cross-reference compounds in another knowledgebase. We determined the cross-references between KEGG and MetaCyc as well as KEGG and Reactome using the kegg-pull Python package (Huckvale and Moseley 2023). We determined the MetaCyc and Reactome cross-references using MetaCyc’s web API (Caspi *et al*. 2020). We trained a model on all the entries in the dataset to make the predictions, predicting on the compounds in the dataset that have a known cross reference in another, resulting in 9193 cross-reference pairs. We arranged the pairs such that the first compound in the pair was one of the compounds in the training set i.e. with known pathway annotations and the second was its cross-reference that may or may not have known pathway annotations in the other knowledgebase. We used the trained models to predict on the training set compounds and then on their cross-references such that the MCC of both can be compared. We additionally counted how many of the cross-reference pairs have both compounds with identical atom color counts.

### Standardizing the dataset

To create a standardized version of the KEGG+Reactome+MetaCyc dataset, we processed the molfiles using the obabel (Open Babel) command line tool (O’Boyle *et al*. 2011). This involved converting the molfiles to InChI format and then back into molfiles, which canonicalizes the chemical representation, i.e. selects a specific tautomeric/resonance form. We created the standardized dataset with the same steps detailed but using the standardized molfiles. We performed the CV, cross-knowledgebase evaluation, and cross-reference analysis detailed above as well.

### Experimenting with atom and bond stereo

The md-harmonize package has the option to specify the atom stereo or bond stereo in the atom colors (Jin and Moseley 2023). Turning the atom or bond stereo off means that these details are not included in the compound feature vectors and therefore the pathway feature vectors, which can impact model performance and generalizability. By default, both atom stereo and bond stereo are turned on for the initial non-standardized dataset and the derived standardized dataset. To determine the impact of atom and bond stereo, we made three more datasets derived from the standardized dataset i.e. one with atom stereo turned on and bond stereo turned off, one with atom stereo turned off and bond stereo turned on, and one with both atom stereo and bond stereo turned off. We performed the CV analysis as well as the cross-reference analysis of each of these three datasets as well.

### Hardware and software used

The hardware used for this work included compute nodes with up to 2 terabytes (TB) of random-access memory (RAM) and central processing units (CPUs) of 3.8 gigahertz (GHz) of processing speed. The CPU was an ‘Intel (R) Xeon (R) Platinum 8480CL’. The graphic processing unit (GPU) was an ‘NVIDIA H100 80 GB HBM3' with 81.56 gigabytes (GB) of GPU RAM in terms of the 1000MB per GB definition. 8 GPUs and 8 cores were used to speed up hyperparameter tuning and CV analysis using multiprocessing where a CV iteration was performed within one of 8 processes at a time.

Table 2 details the computational resources used when training a model on each variant of the dataset. We observe maximal CPU and GPU utilization percentage due to the efficient batching method developed by Huckvale and Moseley (Huckvale and Moseley 2024e) that performs all the batching GPU-side as compared to using multi-processing to perform the batching CPU-side as is done with traditional deep learning batching methods. This 20-fold efficiency is necessary for a dataset of this size, especially when combining all three knowledgebases together. From Table 2, we see that the non-standardized dataset took the most time to train and used the most RAM and GPU RAM.

**Table 2.**
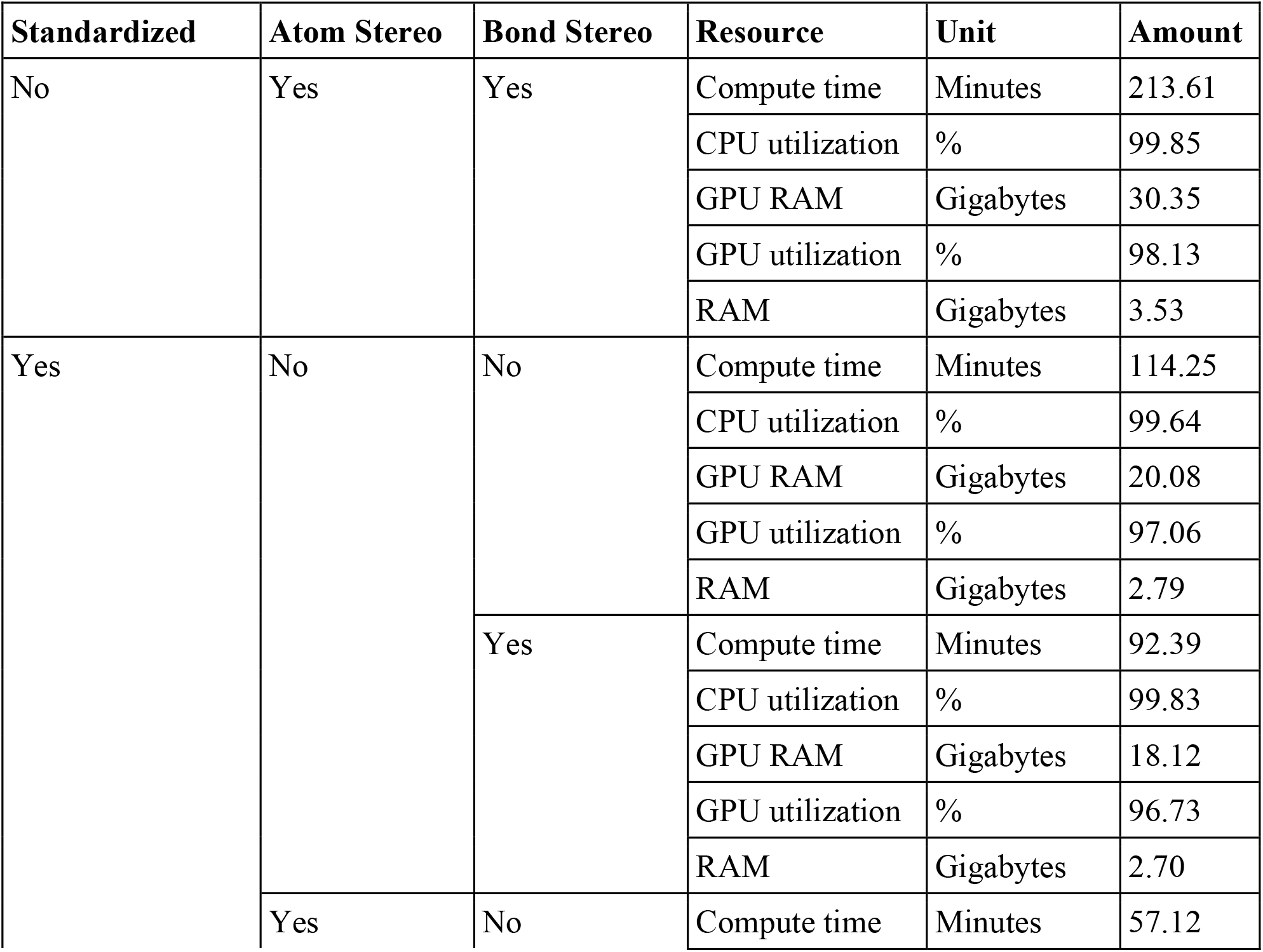
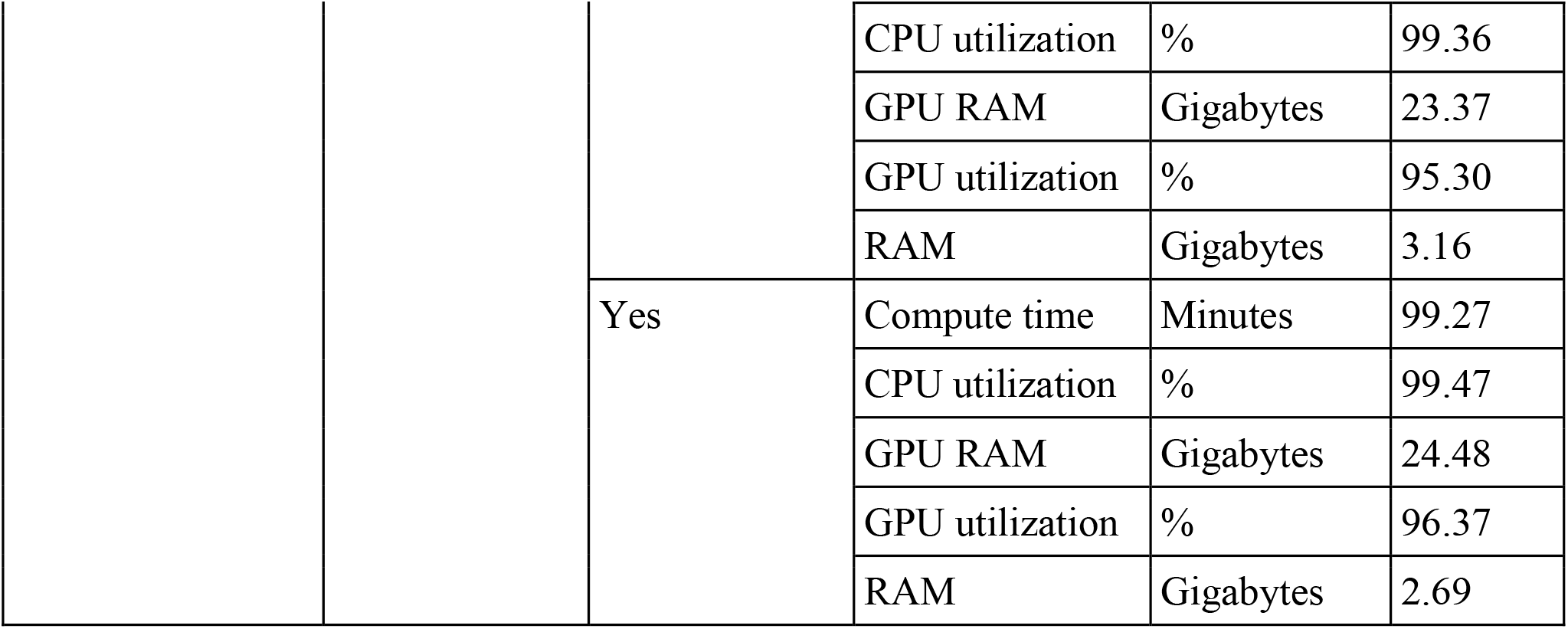
Computational resource usage when training a model on each variant of the dataset.

All code for this work was written in major version 3 of the Python programming language (Rossum and Drake 2009). Data processing and storage were conducted using the Pandas (The pandas development team 2020), NumPy (Harris *et al*. 2020), and H5Py (Collette 2013) packages. Models were constructed and trained using the PyTorch Lightning (Falcon *et al*. 2020) package built upon the PyTorch (Paszke *et al*. 2019) package, gradient descent performed using the Adam optimization algorithm (Kingma and Ba 2014). The stratified train test splits were computed using the Sci-Kit Learn (Pedregosa *et al*. 2012) package. Results were initially stored in an SQL database (Chamberlin 2009) using the DuckDB (Raasveldt and Mühleisen 2019) package. Results were processed and visualized using Jupyter Notebooks (Kluyver *et al*. 2016), the Seaborn package (Waskom 2021) built upon the MatPlotLib (Hunter 2007) package, and the Tableau business intelligence application (Salesforce 2024). Model training and testing were profiled for GPU and CPU utilization using the gpu_tracker package (Huckvale and Moseley 2024f).

## Results

### Non-standardized dataset

#### Model performance

Table 3 includes the MCC from the CV analysis of the models trained on the datasets from the individual KEGG, Reactome, and MetaCyc knowledgebases compared to the KEGG+Reactome+MetaCyc dataset. As demonstrated by prior studies, KEGG by itself has comparable performance to MetaCyc while Reactome significantly exceeds KEGG and MetaCyc. When training a model on all three knowledgebases, the overall mean and median MCC is between that of Reactome and the comparable MCC of KEGG and MetaCyc. However, the standard deviation is roughly half, indicating a large increase in robustness of the models.

**Table 3.** Overall MCC of the individual knowledgebases compared to the combined dataset.

| Dataset | Mean MCC | Median MCC | Standard deviation | Reference |
| --- | --- | --- | --- | --- |
| KEGG+Reactome+MetaCyc | 0.8725 | 0.8737 | 0.0064 | Current study |
| **KEGG** | 0.847 | 0.848 | 0.0098 | (Huckvale and Moseley 2024e) |
| **Reactome** | 0.916 | 0.919 | 0.0149 | (Huckvale and Moseley 2025) |
| **MetaCyc** | 0.8446 | 0.8454 | 0.0101 | (Huckvale and Moseley 2024d) |

Table 4 provides the MCC when predicting the pathways of each knowledgebase using a model trained on the corresponding knowledgebase individually compared to a model trained on the KEGG+Reactome+MetaCyc dataset. When not standardizing the dataset, training on the compounds and pathways of all three knowledgebases increases the average and median MCC when predicting on the pathways in Reactome and MetaCyc and decreases their standard deviations as well. However, the mean and median MCC of KEGG pathways decreases and its standard deviation increases as compared to training on the KEGG data alone, indicating the addition of confusion when predicting KEGG pathways.

**Table 4.**
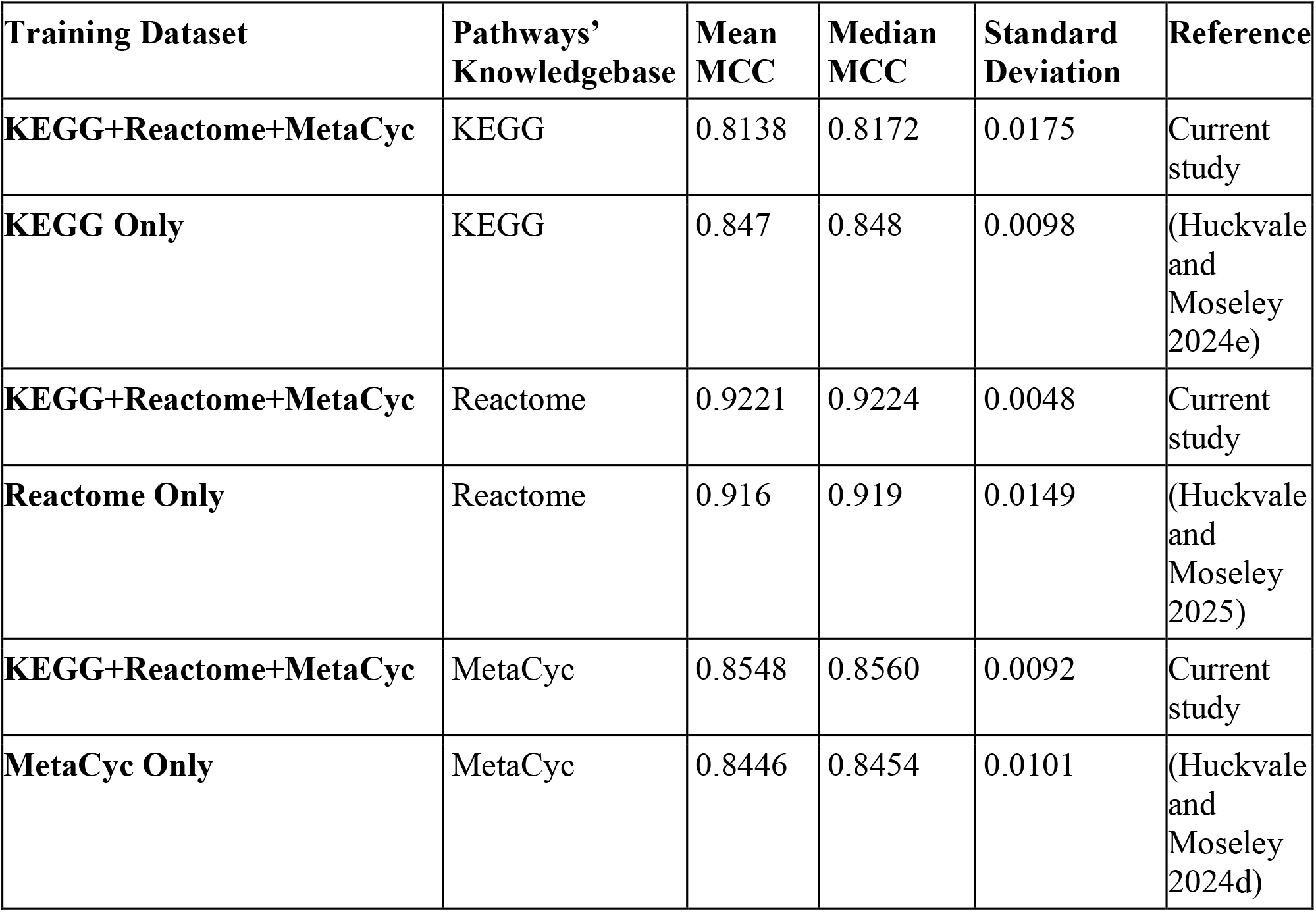
Per-knowledgebase performance when trained on each knowledgebase individually compared to when trained on the combined dataset.

Table 5 shows the MCC when training a model on the entries in one knowledgebase and predicting the pathways of the other two knowledgebases. Regardless of the combination, when training a model on the pathways of one knowledgebase and predicting on the pathways of another knowledgebase, the MCC is very low. Thus, it is very important that the model is trained on the pathways being predicted.

**Table 5.** Cross-knowledgebase evaluation for the non-standardized data.

| Training Knowledgebase | Test Knowledgebase | MCC |
| --- | --- | --- |
| KEGG | Reactome | 0.1196 |
| KEGG | MetaCyc | 0.0455 |
| Reactome | KEGG | 0.1996 |
| Reactome | MetaCyc | 0.1282 |
| MetaCyc | KEGG | 0.2312 |
| MetaCyc | Reactome | 0.2067 |

#### Cross references analysis

Table 6 shows the MCC when using the model trained on all entries to predict the pathways of the training set compounds compared to that of their cross references, which may or may not have been in the training set. It also shows the number of cross-reference pairs that had identical atom color counts. There were 9193 compounds in the training set that had known cross references and out of those, only 1 had identical atom color counts. We see an MCC difference of 0.2687 when predicting on the cross references as compared to the compounds in the training set.

**Table 6.** Comparing the MCC of the compounds in the dataset to that of their cross-references.

| Training set compounds MCC | Cross reference MCC | MCC Difference | #Pairs with identical atom colors |
| --- | --- | --- | --- |
| 0.9129 | 0.6442 | 0.2687 | 1 |

### Standardized dataset

#### Model performance

Table 7 compares the MCC from the CV analysis of the standardized dataset to the non-standardized dataset. Standardizing the chemical representations prior to constructing the atom color features results in a higher MCC and a much lower standard deviation.

**Table 7.** Comparing MCC of the standardized dataset to the non-standardized dataset.

| Standardized | Mean Score | Median Score | Standard Deviation |
| --- | --- | --- | --- |
| Yes | 0.9036 | 0.9040 | 0.0033 |
| No | 0.8725 | 0.8737 | 0.0064 |

Figure 1 provides the same results as Table 7, additionally displaying the distribution of the MCCs across the CV iterations. Both MCC distributions are unimodal. But the MCC distribution for the standardized dataset has a higher center and is less dispersed and has less positive skew.

**Figure 1.**
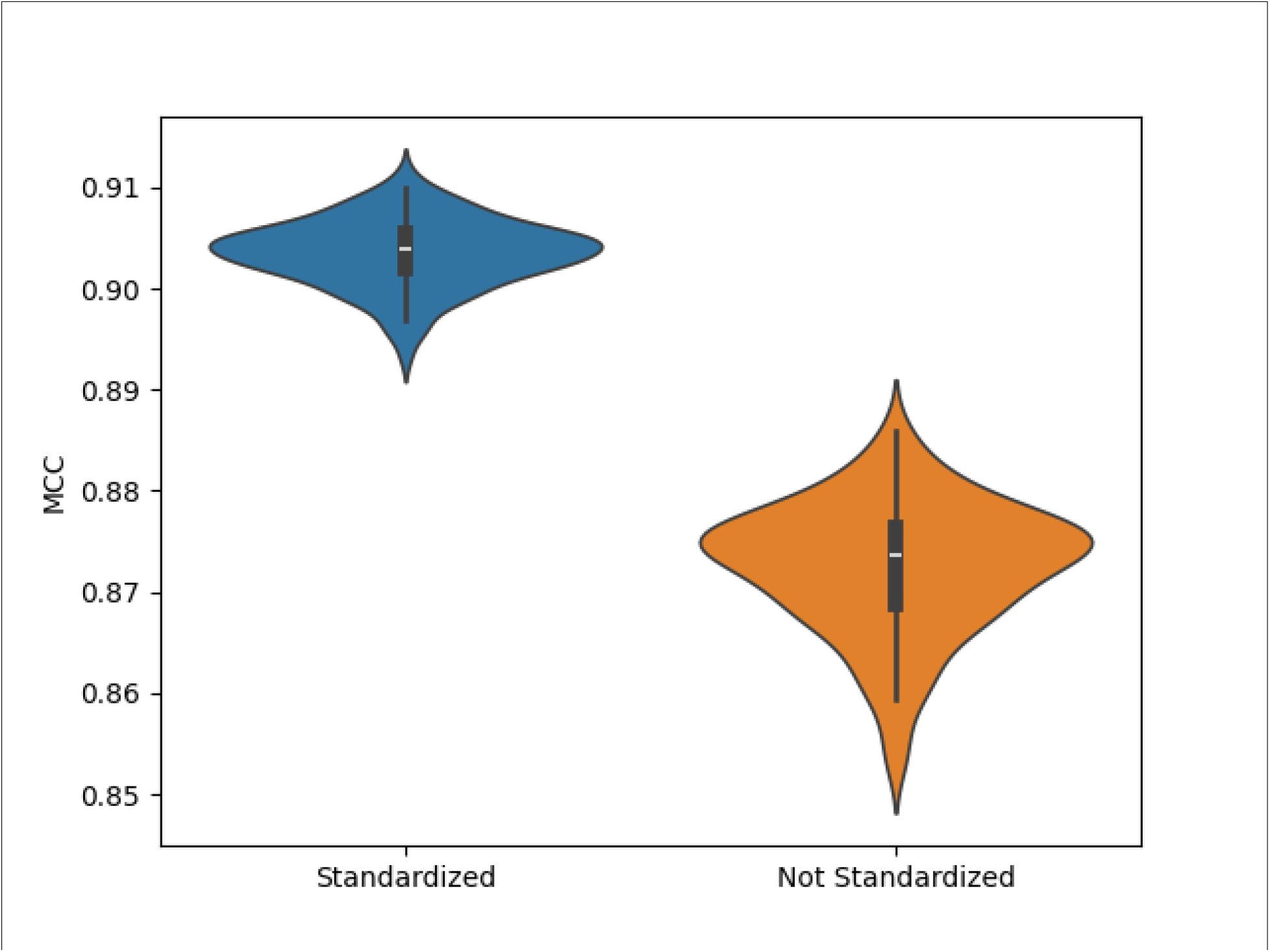
Violin plot of the distribution of the MCC of the standardized and non-standardized datasets across 100 CV iterations.

Table 8 compares the MCC of models trained on the datasets of the individual knowledgebases (which were not standardized) to that of the non-standardized KEGG+Reactome+MetaCyc dataset and the standardized version. The MCC when predicting the pathways of each knowledge is shown for each dataset. Training on the pathways of other knowledgebases increased the MCC and lowered the standard deviation for Reactome and MetaCyc pathways. Standardizing the dataset further increased the MCC and lowered the standard deviation for both. While KEGG pathways initially predicted more poorly when training on all three unstandardized knowledgebases as opposed to training on just KEGG, standardizing the dataset did result in the MCC for KEGG pathways increasing and the standard deviation decreasing when training on other knowledgebases. However, this comparison cannot determine if the improvement is coming from the standardization, the combined dataset, or both.

**Table 8.**
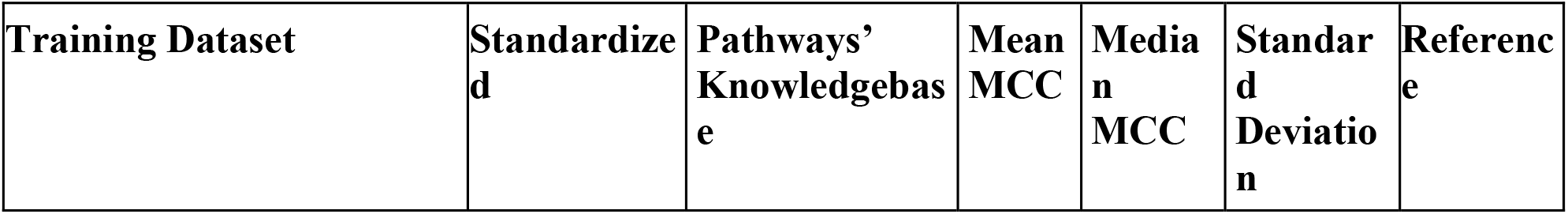
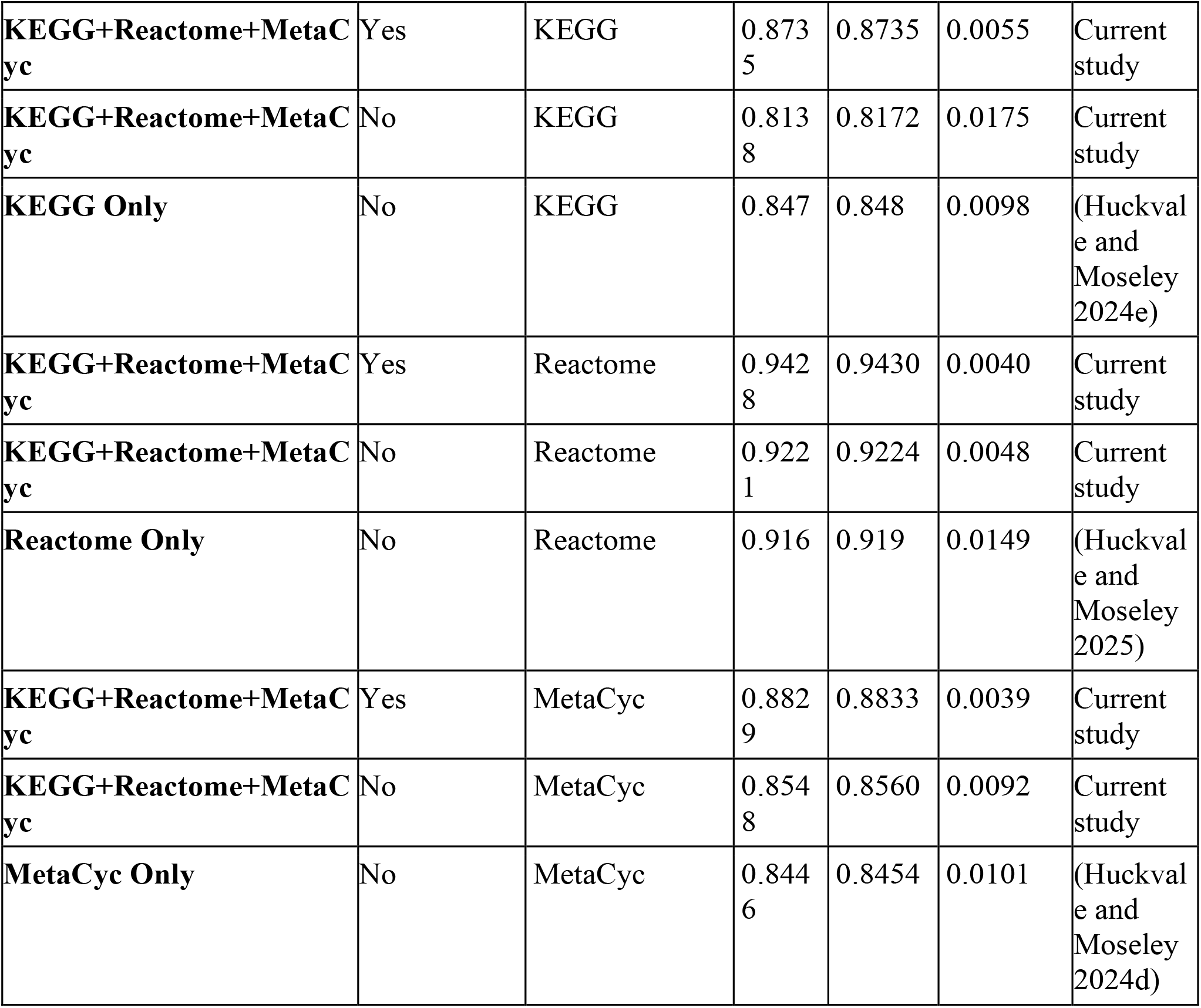
Comparing the MCC when predicting pathways of each knowledgebase using a model trained on the individual knowledgebase, the non-standardized dataset, and the standardized dataset.

Table 9 contains the same results as Table 5 but for the standardized dataset. Standardizing the datasets for knowledgebase prior to predicting on the pathways of another knowledgebase results in better performance than before, but still relatively low. Again, these results highlight how very important that the model is trained on the pathways being predicted.

**Table 9.**
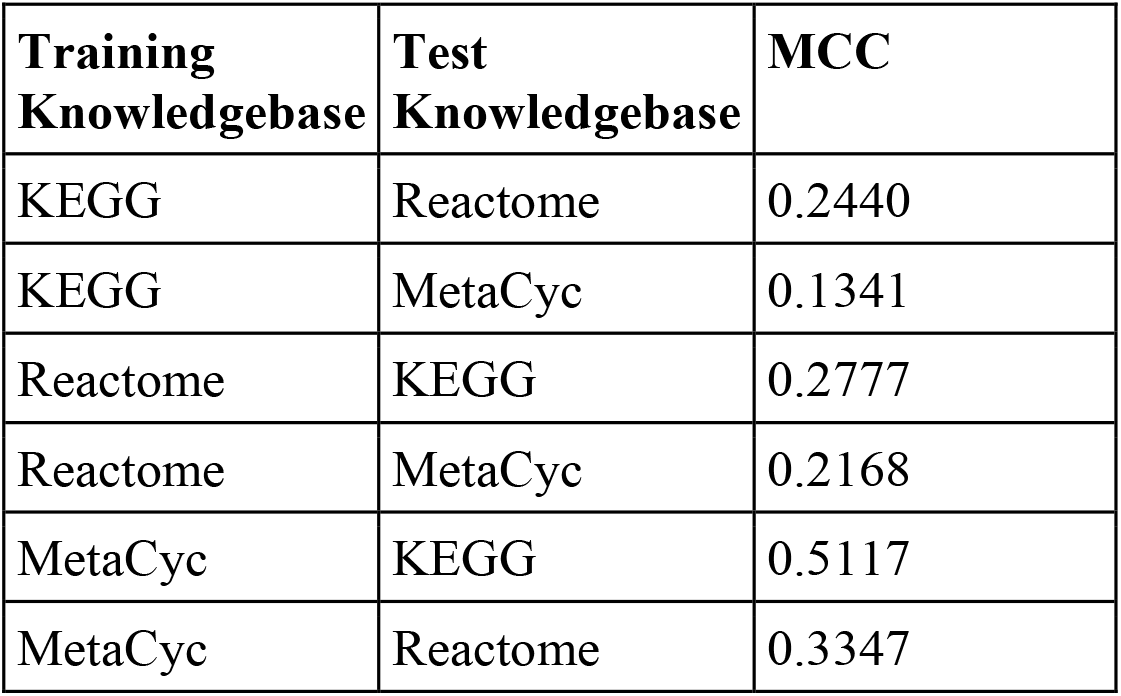
Cross-knowledgebase evaluation for the standardized data.

#### Cross reference analysis

Table 10 compares the data and prediction consistency when standardizing the data and when not standardizing the data. The standardized dataset results in increased MCCs when predicting on the compounds with known pathway annotations. More importantly, it greatly increases the MCC of the cross references. And the difference between the training set compounds and their cross references is much smaller, significantly improving consistency. This is largely attributed to by the increased number of cross reference pairs that have identical atom color counts after standardizing, increasing from 1 out of 9193 to 7234 out of 9193 ≈ 78.7%.

**Table 10.** Comparing the prediction and data consistency when standardizing and when not standardizing.

| Standardized | Training set compounds MCC | Cross reference MCC | MCC Difference | #Pairs with identical atom colors |
| --- | --- | --- | --- | --- |
| No | 0.9129 | 0.6442 | 0.2687 | 1 |
| Yes | 0.9623 | 0.9239 | 0.0384 | 7234 |

#### Atom stereo and bond stereo inclusion

**Error! Reference source not found**. shows the number of training set compounds with identical atom colors to their cross references and how these numbers are impacted by whether atom stereo or bond stereo is turned on or off when generating atom colors. While standardizing resulted in many more compounds having identical atom colors, removing the atom and bond stereo details from the atom colors resulted in even more identical compounds, making the compound representation more consistent. See Table S2 for the same counts but for SMILES standardization instead of InChI. SMILES standardization resulted in less consistency, justifying the use of InChI for this use case. The prediction consistency likewise increased, since we see a smaller MCC difference between the training set compounds and their cross-references. However, the best cross-reference MCC of 0.9239 was observed when using InChI standardization with both atom and bond stereo on.

**Table 11.** Comparison of prediction and data consistency for the four different combinations of atom and bond stereo inclusion.

| Standardized | Atom Stereo | Bond Stereo | Training set compounds MCC | Cross reference MCC | MCC Difference | #Pairs with identical atom colors |
| --- | --- | --- | --- | --- | --- | --- |
| No | On | On | 0.9129 | 0.6442 | 0.2687 | 1 |
| Yes | On | On | 0.9623 | 0.9239 | 0.0384 | 7234 |
|  |  | Off | 0.8965 | 0.8679 | 0.0287 | 7465 |
|  | Off | On | 0.9320 | 0.9164 | 0.0155 | 8510 |
|  |  | Off | 0.8871 | 0.8773 | 0.0098 | 8762 |

Table 12 shows the results of the CV analysis of the four combinations of atom and bond stereo being turned on and off for the standardized dataset. While excluding atom stereo and bond stereo details results in greater prediction and data consistency between cross references, it reduces the overall model performance. See Table S3 for these results for all metrics in addition to MCC and for all combinations of whether the data is standardized and whether atom or bond stereo is turned on or off. Again, the best performance was observed with atom and bond stereo on, producing a mean MCC of 0.9036 ± 0.0033.

**Table 12.** CV analysis of the four combinations of atom and bond stereo inclusion.

| Atom Stereo | Bond Stereo | Mean MCC | Median MCC | Standard Deviation |
| --- | --- | --- | --- | --- |
| On | On | 0.9036 | 0.9040 | 0.0033 |
| On | Off | 0.8707 | 0.8713 | 0.0063 |
| Off | On | 0.8985 | 0.8984 | 0.0047 |
| Off | Off | 0.8840 | 0.8859 | 0.0089 |

## Discussion

We combined the KEGG, Reactome, and MetaCyc knowledgebases together to create a single combined dataset with 16,640 unique compound feature vectors, 8,195 unique pathway feature vectors, and 50,127,958 compound-pathway entries. With the new combined dataset, the robustness of the resulting models improves to a mean MCC of 0.9036 ± 0.0033, with the standard deviation less than one third of all prior published results. These are the best results published so far and are far better than older multiclassifiers and one-vs-rest binary classifiers that all had prediction performance on average below an MCC of 0.8. Moreover, all models prior to September 21, 2024 only predicted 11 or 12 level 2 KEGG metabolic pathways versus over 8000 pathways predicted by the extreme classification model presented here.

The extreme classification model performance when predicting on Reactome pathways and MetaCyc pathways additionally improves, indicating transfer learning across the knowledgebases. However, KEGG pathway prediction performance decreases. The lower performance and robustness of KEGG pathways when trained along with the other two knowledgebases is caused by confusion introduced by inconsistent chemical representations between the knowledgebases. One might conclude that it is advisable to use a model trained on KEGG pathways only when predicting KEGG pathways. However, standardizing the chemical representations with InChI canonicalization evidently corrects and/or compensates for this discrepancy when training a model on all three knowledgebases, with KEGG pathway prediction performance improving. Chemical representation standardization further improves Reactome and MetaCyc as well. These results taken together indicate that standardizing the chemical representation of compounds significantly improves both model performance and robustness by enabling additional transfer learning between knowledgebase pathways and/or preventing confusion, depending on your perspective.

Moreover, our cross-reference analyses demonstrate the high inconsistency in chemical representations across knowledgebases with only 1 out of 9,193 cross-reference pairs having identical atom coloring feature vectors. After chemical representation standardization, consistency across knowledgebases increases dramatically to 7,234 out of 9,193 ≈ 78.7%. By removing these inconsistencies in chemical representation, the drop in MCC for the cross-references goes from 0.2687 without standardization to 0.0384 with standardization. Also, the standardized cross-reference MCC is 0.9239, which is the highest performance. Thus, the resulting models are more generalizable when predicting on compound entries outside of the training data while also maintaining high prediction performance. Therefore, it is essential to standardize the data prior to predicting metabolic pathway involvement. To our knowledge, investigation into data engineering techniques to maximize model generalizability across different knowledgebases with different chemical representations has not been previously published.

Also, the method of standardization matters. Using SMILES for standardization was not as useful as using InChI (Table S2). Also, the three knowledgebases have their own standardizations. However, different standardizations can have different tautomeric, resonance, and ionization preferences in chemical representation, which is illustrated by the poor performance when the three knowledgebases are combined without a separate standardization step. Likewise, PubChem’s standardization has a 60% inconsistency with InChI canonicalization (Hähnke, Kim and Bolton 2018). Again, this all supports the use of a single chemical representation standardization method prior to training and predicting metabolic pathway involvement.

While the extreme classification models presented here have significantly higher performance than all prior published results, there are still limitations. While these models generalize to novel chemical representation of compounds, they do not generalize well to novel pathways. This is evident by the poor performance when building models trained on one knowledgebase and then predicting the pathways of another knowledgebase. When predicting metabolic pathways on novel compounds, we recommend predicting only pathways that the model was trained on. Further research is required to determine a way to generalize on novel pathways.

Use of a multi-layer perceptron (MLP) model facilitates the cross-join technique for extreme classification since input features in a graph format cannot be cross-joined to vectorized pathway features. If a graph2vec approach is used to vectorize the graph features prior to the cross-join to the pathway feature vectors, there are still GPU memory limitations for a dataset of this size, which we have directly tested. If the batch size is decreased in order to process a smaller number of compounds and prevent the graph neural network from overloading GPU memory, the batch size would be too small for the model to train in a reasonable amount of time. One could batch the compounds alone, perform graph2vec, and then cross-join to the pathway features, but current batching techniques as provided by deep learning libraries such as PyTorch are done CPU side with multiprocessing where additional time is consumed transferring data in between processes. However, we needed to create our own batching mechanisms performed entirely on the GPU side, and in the same process, in order to practically train a model on a dataset of this size. Our custom batching method (data loading method) improved GPU utilization 20-fold, making the current model training, testing, and evaluation practical on a dataset with over 50,000,000 entries. Special batching techniques would need to be developed to allow the use of a graph neural network followed by a cross-join of the vectorized compound representations. Such batching techniques are non-trivial to implement for graph data. However, here we demonstrate excellent performance using vector representations with atom color features and a MLP model. Results may be improved if a batching technique with graph representations is implemented and that is efficiently performed on the GPU side, making the batching practical for a dataset of this size.

## Conclusions

The KEGG+Reactome+MetaCyc dataset contains 16,640 uniquely represented compounds, 8,195 uniquely represented pathways, and 50,127,958 compound-pathway entries. Our extreme classification MLP models can predict 8,195 pathways with a mean MCC 0.9036 ± 0.0033, the best results published so far. The chemical representation standardization using InChI canonicalization significantly improves model performance and is essential for generalizability when predicting metabolic pathway involvement on novel compounds that would otherwise have an inconsistent chemical representation with the model. At this time, we recommend predicting the same pathways that the model was trained to predict. While there may be further improvement by using graph neural network methods, this requires custom batching techniques to be developed and implemented for training and testing to be practical on a dataset of this size (tens of millions of entries).

## Acknowledgements

We thank the University of Kentucky Institute for Biomedical Informatics and National Science Foundation Grant Number 1626364 for their support and associated computing resources.

---

**Table 1.** MCC of the dataset and model introduced by Baranwal et al after 50 CV iterations.

| Mean MCC | Standard deviation |
| --- | --- |
| 0.7642 | 0.0137 |

**Table 2.** Number of identical atom color counts between training set compounds and their cross-references when standardizing the data by converting to SMILES format.

|  | Bond Stereo On | Bond Stereo Off |
| --- | --- | --- |
| Atom Stereo On | 2066 | 2675 |
| Atom Stereo Off | 6031 | 7495 |

**Table 3.** CV analysis for all metrics and all combinations of standardization, atom stereo, and bond stereo

| Atom Stereo | Bond Stereo | Standardized | Metric | Mean Score | Median Score | Standard Deviation |
| --- | --- | --- | --- | --- | --- | --- |
| atom-stereo | bond-stereo | not-standardized | Accuracy | 0.9984 | 0.9985 | 0.0001 |
| atom-stereo | bond-stereo | not-standardized | F1 score | 0.8696 | 0.8708 | 0.0070 |
| atom-stereo | bond-stereo | not-standardized | MCC | 0.8725 | 0.8737 | 0.0064 |
| atom-stereo | bond-stereo | not-standardized | Precision | 0.7975 | 0.7986 | 0.0129 |
| atom-stereo | bond-stereo | not-standardized | Recall | 0.9562 | 0.9566 | 0.0035 |
| atom-stereo | bond-stereo | standardized | Accuracy | 0.9989 | 0.9989 | 0.0000 |
| atom-stereo | bond-stereo | standardized | F1 score | 0.9031 | 0.9035 | 0.0036 |
| atom-stereo | bond-stereo | standardized | MCC | 0.9036 | 0.9040 | 0.0033 |
| atom-stereo | bond-stereo | standardized | Precision | 0.8629 | 0.8636 | 0.0092 |
| atom-stereo | bond-stereo | standardized | Recall | 0.9473 | 0.9476 | 0.0039 |
| atom-stereo | no-bond-stereo | standardized | Accuracy | 0.9984 | 0.9985 | 0.0001 |
| atom-stereo | no-bond-stereo | standardized | F1 score | 0.8675 | 0.8682 | 0.0070 |
| atom-stereo | no-bond-stereo | standardized | MCC | 0.8707 | 0.8713 | 0.0063 |
| atom-stereo | no-bond-stereo | standardized | Precision | 0.7929 | 0.7942 | 0.0139 |
| atom-stereo | no-bond-stereo | standardized | Recall | 0.9578 | 0.9580 | 0.0038 |
| no-atom-stereo | bond-stereo | standardized | Accuracy | 0.9988 | 0.9988 | 0.0001 |
| no-atom-stereo | bond-stereo | standardized | F1 score | 0.8974 | 0.8974 | 0.0051 |
| no-atom-stereo | bond-stereo | standardized | MCC | 0.8985 | 0.8984 | 0.0047 |
| no-atom-stereo | bond-stereo | standardized | Precision | 0.8467 | 0.8471 | 0.0120 |
| no-atom-stereo | bond-stereo | standardized | Recall | 0.9547 | 0.9547 | 0.0042 |
| no-atom-stereo | no-bond-stereo | standardized | Accuracy | 0.9986 | 0.9986 | 0.0001 |
| no-atom-stereo | no-bond-stereo | standardized | F1 score | 0.8817 | 0.8839 | 0.0099 |
| no-atom-stereo | no-bond-stereo | standardized | MCC | 0.8840 | 0.8859 | 0.0089 |
| no-atom-stereo | no-bond-stereo | standardized | Precision | 0.8159 | 0.8196 | 0.0202 |
| no-atom-stereo | no-bond-stereo | standardized | Recall | 0.9596 | 0.9603 | 0.0054 |

